# Occurrence of “under-the-radar” antibiotic resistance in anthropogenically affected produce

**DOI:** 10.1101/2024.09.26.615221

**Authors:** Chagai Davidovich, Kseniia Erokhina, Chhedi Lal Gupta, Yong-Guan Zhu, Jian-Qiang Su, Steven P. Djordjevic, Ethan R. Wyrsch, Shlomo Blum, Eddie Cytryn

## Abstract

With global climate change, treated-wastewater (TWW) irrigation and manure amendment are becoming increasingly important in sustainable agriculture in water- and nutrient-stressed regions. Yet, these practices can potentially disseminate pathogens and antimicrobial resistance (AMR) determinants to crops, resulting in serious health risks to humans through the food chain. Previous studies demonstrated that pathogen and AMR indicators from wastewater and manure survive poorly in the environment, suggesting that ecological barriers prevent their dissemination. However, we recently found that these elements can persist below detection levels in low quality TWW-irrigated soil, and potentially proliferate under favorable conditions. This “under the radar” phenomenon was further investigated here, in TWW irrigated- and poultry litter-amended lettuce plants, using an enrichment platform that resembles gut conditions, and an analytical approach that combined molecular and cultivation-based techniques. Enrichment uncovered clinically-relevant multidrug resistant pathogen indicators and a myriad of antibiotic resistance genes in the litter amended and TWW-irrigated lettuce that were not detected by direct analyses, or in the enriched freshwater irrigated samples. Selected resistant *E. coli* isolates were capable of horizontally transferring plasmids carrying multiple resistance genes to a susceptible strain. Overall, our study underlines the hidden risks of under-the-radar pathogen and AMR determinants in anthropogenically affected agroenvironments, providing a novel platform to improve quantitative microbial risk assessment models in the future.

## Introduction

Antimicrobial resistance (AMR) is one of the most significant global threats to public health in the 21st century^1^, and containment and mitigation of AMR necessitates comprehensive understanding of its emergence and dissemination routes. While the clinical implications of AMR are widely recognized, there is an increasing awareness of the environmental dimension of this problem^2,3^. The “One-Health” concept underlines the need to examine AMR holistically, emphasizing relationships between humans and animals, and specifically the food chain that links the two^4^. Animal husbandry and aquaculture have drawn attention as potential hotspots for the development and dissemination of antibiotic resistance due to the extensive use of antibiotics^3^. Moreover, agronomic practices aimed at sustaining crop productivity, such as wastewater irrigation and manure amendment, can introduce a myriad of antibiotic-resistant bacteria (ARB) and antibiotic resistance genes (ARGs) originating from human and animal wastes, into soil-crop ecosystems, thus contributing to AMR dissemination^5^.

Molecular and culture-based methods are the “gold standards” for quantifying ARB and ARGs in agroecosystems^6,7^. However, it is important to recognize the limits of detection (LOD) of individual methods, and realize that failure to detect a specific ARB/ARG in a targeted environment does not mean that it is not present^8^. This realization is crucial because under nutrient-rich conditions such as in the gut following ingestion of produce, resistant pathogens can proliferate, and persistent bacteria harboring mobile genetic elements (MGEs) containing ARGs can be horizontally transferred to native strains^9,10^.

Enrichment methods are traditionally applied in diagnostic microbiology laboratories to enhance isolation of specific bacterial pathogens such as *Salmonella* spp. and *Listeria monocytogenes* in clinical or food samples^11,12^, but enrichment can also be applied to environmental surveillance studies that focus on multiple strains and not just one specific pathogens. For example, by applying a nonselective enrichment step, Blau *et. al.* determined that tetracycline resistant bacteria from commercial produce were capable of transferring their resistance to bacterial pathogen indicators^13^. In a separate study conducted in our lab, short-term enrichment of freshwater and treated wastewater (TWW) irrigated soils in a copiotrophic medium facilitated detection of clinically relevant ARB (including cephalosporin- and carbapenem-resistant *Enterobacteriaceae*) and associated ARGs in TWW irrigated soils that were not present in the corresponding freshwater irrigated soils^14^. Collectively, these studies highlight the fact that although enrichment causes bias, and enriched microbiomes are not representative of native bacterial communities, it can potentially uncover ESKAPE pathogens^15^ and ARG-harboring mobile elements, and thus can be valuable for epidemiological surveillance of environmental samples.

This study evaluated the contribution of TWW irrigation and poultry-litter amendment to the dissemination of bacterial pathogen and ARG indicators of clinical concern, in the rhizosphere and phyllosphere of lettuce, as a model for crop production. Rhizosphere and phyllosphere samples were analyzed both directly and following enrichment under aerobic and anaerobic conditions, mimicking conditions in the human gut. We incorporated a methodological approach that combined culture-based techniques that targeted total and extended spectrum β-lactamase producing *E. coli*^16–18^, with molecular methods, including high throughput qPCR, 16S rRNA amplicon sequencing, and shotgun metagenomic analysis. Enrichment revealed clinically relevant ARB and ARGs in TWW-irrigated and litter-amended produce that were not observed by direct analyses or in enriched freshwater irrigated samples. These “under the radar” determinants could have substantial, yet overlooked, epidemiological importance, underlining the potential of environmental enrichment platforms in future surveillance and microbial risk assessment programs.

## Results

### Enrichment facilitates significant shifts in bacterial diversity and composition

We evaluated bacterial diversity and community structure of native and enriched freshwater, litter-amended, and TWW-irrigated lettuce rhizosphere and phyllosphere samples using 16S rRNA amplicon sequencing. Rhizosphere bacterial diversity was significantly higher than that of the phyllosphere, and native samples were significantly more diverse than the enriched phyllosphere and rhizosphere ones (Wilcoxon rank-sum test p<0.00001; Fig. 1A; supplementary Fig. 1A). Enrichment induced significant shifts in the bacterial community structure of both rhizosphere and phyllosphere samples (permutational multivariate analysis of variance (PERMANOVA) P.adj < 0.001). Distinct bacterial communities were observed in both native and enriched freshwater, litter-amended and TWW-irrigated rhizosphere samples. In contrast, in phyllosphere samples significant differences were observed in the communities of the three treatments in the enriched samples, but not in the native ones (PERMANOVA p.adj<0.05. Fig. 1B). A positive correlation between α-diversity and the level of “anthropogenicity” (TWW irrigation > litter amendment > freshwater irrigation) was found in native and enriched rhizosphere samples but not in the phyllosphere ones, and this effect was stronger in enrichments than in native samples (Wilcoxon rank-sum test p<0.05; Fig. 1C).

**Fig. 1.**
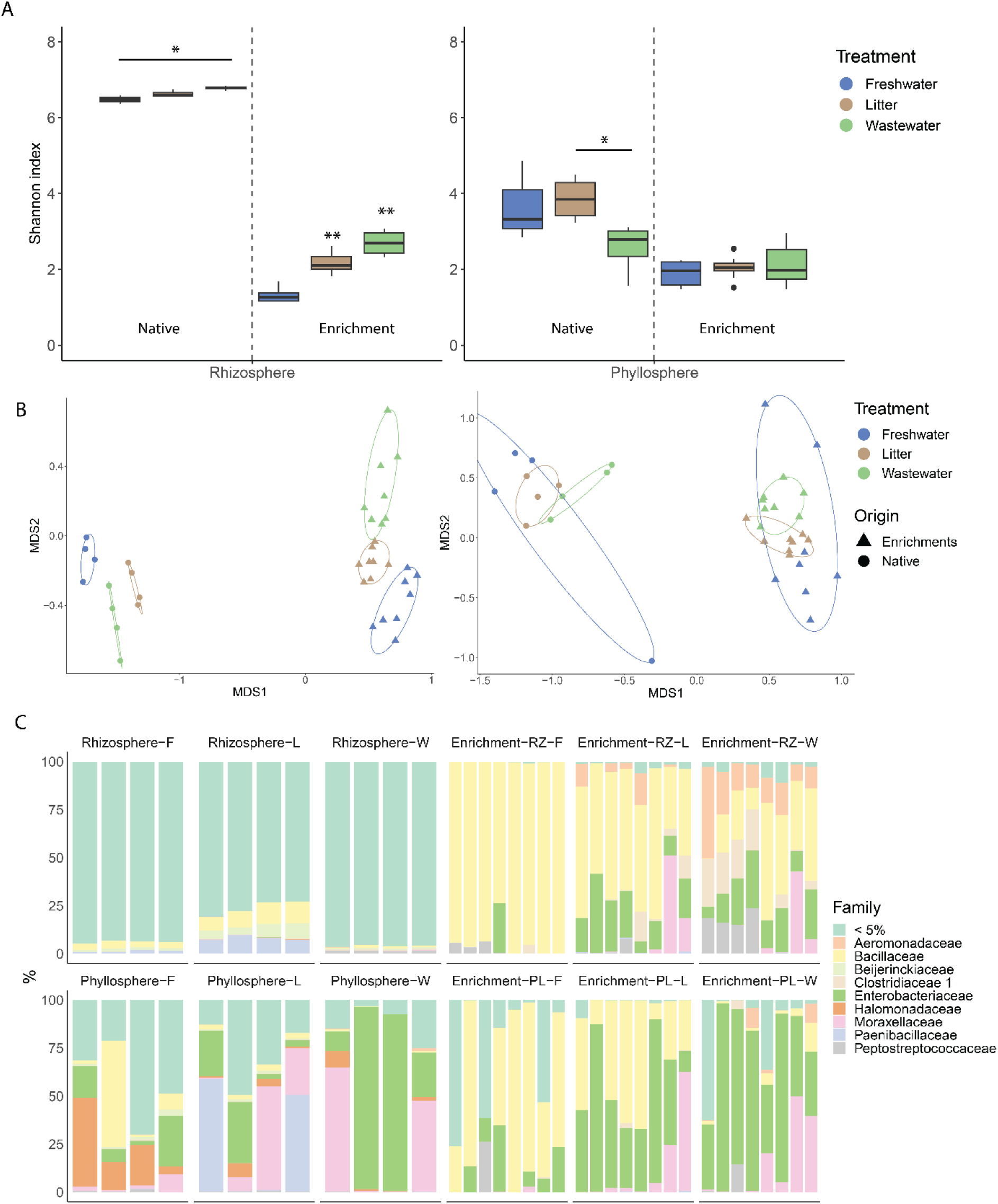
Impact of treatment and enrichment on bacterial diversity and community composition based on analysis of 16S rRNA gene amplicon sequencing data. **A** α-diversity of native and enriched freshwater-irrigated, litter-amended, and TWW-irrigated rhizosphere and phyllosphere samples measured by the Shannon diversity index (pairwise Wilcoxon test * p.adj < 0.05; ** p.adj < 0.005). **B** β-diversity of native and enriched rhizosphere (left) and phyllosphere (right) samples, measured using Bray–Curtis distance matrices, and visualized by NMDS plot analysis (PERMANOVA, p.value < 0.05). **C** Bacterial community composition (family-level) of native and enriched rhizosphere and phyllosphere samples. Aerobic and anaerobic enrichments were combined because redox potential (aerobic vs. anaerobic inoculations) did not significantly impact the overall bacterial community structure. RZ= rhizosphere, PL= phyllosphere, F= freshwater, L= litter, W= TWW. Bars represent biological replicates. <5% indicates families whose relative abundance represents < 5% of the total characterized bacterial community.

The native rhizosphere microbiome was highly diverse, and relatively homogeneous, with the majority of characterized phyla representing less than 5% of the total bacterial community. In contrast, the native phyllosphere microbiome was less diverse, and substantially varied between biological replicates. In both phyllosphere and rhizosphere samples, enrichment resulted in almost complete takeover of families associated with the Pseudomonadota and Bacillota phyla. However, enrichment of freshwater irrigated samples primarily stimulated proliferation of *Bacillaceae*, whereas enrichment of litter-amended and TWW-irrigated samples enhanced the relative abundance of *Aeromonadaceae*, *Moraxelaceae*, and *Enterobacteriaceae*, that are more strongly associated with clinically relevant, opportunistic and obligatory pathogens^19^. This phenomenon was significant both in enriched rhizosphere and phyllosphere microbiomes despite the high variability of the native phyllosphere samples, emphasizing the potential of copiotrophic enrichment to reveal clinically relevant taxa in highly diverse environmental samples (Fig. 1C and supplementary Fig. 1C).

### Clinically relevant bacteria persist in the rhizosphere and phyllosphere below LOD and proliferate following enrichment

We applied differential abundance analysis of the 16S rRNA gene amplicon data to identify taxa that were significantly more abundant in the enriched samples compared to their corresponding native rhizosphere and phyllosphere samples. Analysis showed that bacterial pathogen-associated genera, including *Aeromonas*, *Escherichia-Shigella*, *Acinetobacter*, *Enterococcus* and unclassified *Enterobacteriaceae* were significantly more abundant in enriched anthropogenically impacted rhizosphere TWW-irrigated samples then their native counterparts. Similarly, unclassified *Enterobacteriaceae*, *Aeromonas*, and *Acinetobacter* were significantly more abundant in enriched litter-amended rhizosphere samples than in the corresponding native samples. Most notably, *Escherichia-Shigella* and *Enterococcus* were only detected in the rhizosphere TWW-irrigated samples following enrichment, underlining the capacity of enrichment to detect “under-the-radar” pathogen indicators. Although evident, the anthropogenic footprint of the enriched phyllosphere samples was less distinct than that of the enriched rhizosphere samples, but was still evident in the case of unclassified *Enterobacteriaceae*, which were more abundant in enriched litter-amended phyllosphere samples compared to native litter amended phyllosphere samples (DESeq2 log2FoldChange > 1, padj < 0.01; Fig. 2A and supplementary Fig. 2).

**Fig. 2.**
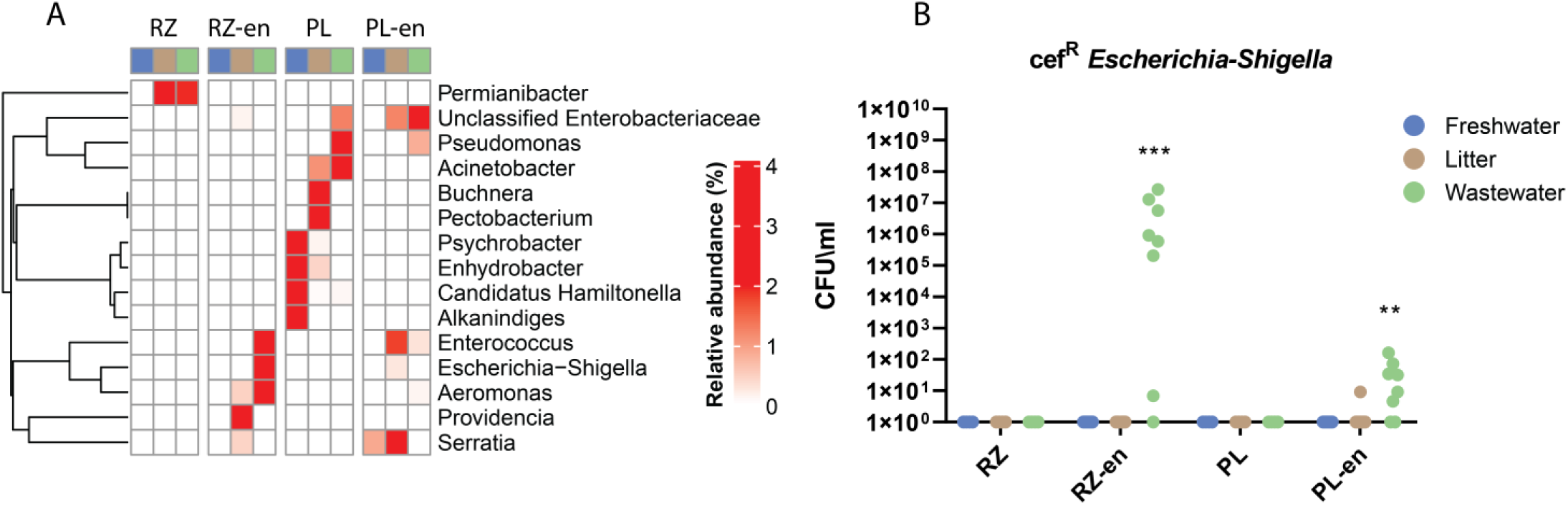
Effect of treatment of differential abundance of taxa in native and enriched rhizosphere and phyllosphere samples. **A** Differential abundance of clinically-relevant genera extrapolated from 16S rRNA gene amplicon sequencing data using DESeq2 (log_2_FoldChange > 1, padj < 0.01). Each square represents the average values of 4 replicates for native samples and 8 replicates for enriched (aerobic and anaerobic) samples. **B** Quantification of cefotaxime resistant *E.coli* (a gold-standard ESBL-producing fecal pathogen indicator) in native and (aerobic and anaerobic) enriched samples. Resistant *E. coli* isolates were detected using a selective CCA agar supplemented with cefotaxime (4 µg/ml).Taxonomy of representative isolates was validated by MALDI-TOF analysis. Wilcoxon test was applied for multiple comparisons *** p-value <0.0001; ** p-value<0.001; cef^R^= cefotaxime resistant.

Complementation of 16S rRNA gene amplicon data with shotgun metagenomic analysis of enriched samples identified several pathogen-associated *Enterobacteriaceae* genera that were substantially more abundant in the anthropogenically-impacted rhizosphere enrichments, including *Salmonella*, *Enterobacter*, and *Klebsiella*, which belong to the unclassified-*Enterobacteriaceae* fraction characteristic of enriched anthropogenically impacted samples background (Supplementary Fig. 5B).

Collectively, bacterial community analyses revealed that enrichment facilitated proliferation of *Bacillus* in all treatments regardless of environmental conditions. Aerobic enrichment resulted in higher abundances of *Pseudomonas* and *Acinetobacter*, whereas *Paraclostridium* proliferated in the anaerobic enrichments, coinciding with results from previous studies^20,21^. As shown, anthropogenic background gave rise to the proliferation of pathogen-associated genera, i.e. *Escherichia-Shigella*, *Salmonella*, *Enterobacter*, *Klebsiella*, *Enterococcus*, *Aeromonas* and *Acinetobacter* (Supplementary table 1).

Concomitant to the molecular analyses, we applied culture-based analysis to evaluate the abundance of total and cefotaxime-resistant fecal coliforms and *E. coli* by plating on Chromocult Coliform Agar (CCA) and subsequently characterizing taxonomic affiliation of colonies using Matrix-assisted laser desorption/ionization time-of-flight mass spectrometry (MALDI-TOF MS). Cefotaxime resistant *E. coli* were not detected in the native TWW-irrigated rhizosphere and phyllosphere samples, however, enrichment caused these bacteria to proliferate in both TWW-irrigated rhizosphere and phyllosphere samples. Similarly, cefotaxime-resistant fecal coliforms were quantified in the enriched litter-amended and TWW-irrigated samples, but not in the respective native samples. In contrast, cefotaxime-resistant *E. coli* and fecal coliforms were not present in any of the freshwater irrigated samples regardless of enrichment (Fig. 2B and supplementary Fig. 3). These results highlight the capacity of enrichment to reveal presence of gold-standard bacterial pathogen and ARB indicators that are below detection limits in direct culture-based and culture-independent analyses.

### Enrichment reveals under-the-radar ARGs and MGEs in anthropogenically impacted rhizosphere and phyllosphere samples

To determine the effects of enrichment and anthropogenic exposure on the scope of ARGs and MGEs and to zoom-in on specific pathogen associated taxa, DNA from the enriched rhizosphere and phyllosphere samples was subjected to shotgun metagenomic sequencing and high-throughput quantitative PCR (HTqPCR). Shotgun metagenomic analysis revealed positive correlations between the level of anthropogenic disturbance (TWW-irrigation > litter amendment > freshwater irrigation), and the abundance and diversity of clinically relevant ARGs and MGEs in the rhizosphere and phyllosphere enrichments. Notable clinically- and plasmid-associated ARGs that were significantly more abundant in the enriched anthropogenically affected samples included β - lactamases such as *bla*_TEM_, *bla*_CTX-M_, *bla*_SHV_, and the carbapenemase *bla*_VIM-1_, as well as pathogen-associated genes, and genes encoding resistance to tetracyclines, sulfonamides, and fluoroquinolones^22,23^ (Fig. 3A and supplementary Fig. 5A).

**Fig. 3.**
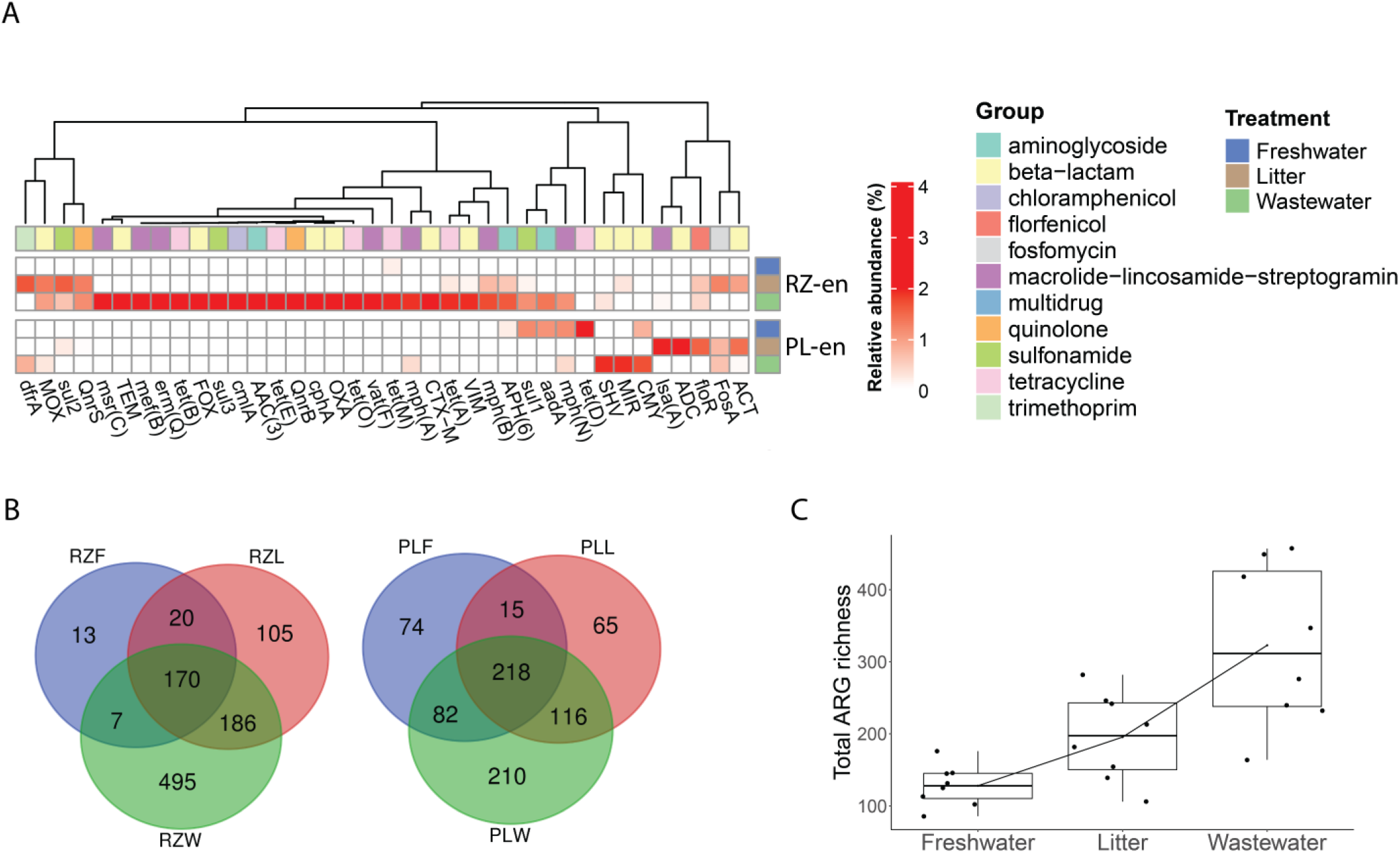
Relative abundance and diversity of ARGs extrapolated from shotgun metagenomes of enriched samples. **A** Average relative abundance (n=4) of differentially abundant ARGs (DESeq2 log_2_FoldChange > 1, padj < 0.01) in enriched rhizosphere and phyllosphere samples, focusing on mobile ARGs (identified by aligning DESeq2 results to the ResFinder database^24^) that were significantly more abundant in anthropogenically-associated enrichments than in freshwater-irrigated samples. **B** Venn diagram showing distinct and shared ARGs between freshwater, litter, and TWW-associated enrichments. **C** ARG richness of enriched rhizosphere and phyllosphere samples as a function of treatment. The line shows the increasing trend in ARG richness between freshwater-irrigated, litter-amended and TWW-irrigated produce (p.adj < 0.05, Wilcoxon test). RZ- rhizosphere; PL-phyllosphere. F- freshwater; L- litter; W-wastewater; en-enrichments.

Similar to the shotgun metagenomic analysis, HTqPCR identified several markers in the anthropogenically-impacted enriched samples that were not present in the freshwater-irrigated ones. Most notably was the proliferation of ARGs conferring resistance to third-generation cephalosporins, fluoroquinolones, and tetracyclines, and of *Enterococci*, *Acinetobacter baumannii*, and *Klebsiella pneumoniae* specific markers. Most of the significant gene markers were exclusively present in the enriched rhizosphere samples. However, 6 out of 34 markers that were found to be significantly more abundant in anthropogenically associated rhizosphere enrichments, were also found to be significant in the enriched litter-amended and TWW-irrigated phyllosphere samples, including species-specific markers for *K. pneumoniae* and *A. baumannii*, and β-lactamases such as *bla*_TEM_ (Wilcoxon test, P < 0.05; supplementary Fig. 4).

The anthropogenic footprint of the enriched samples was clearly evident when observing total ARG richness and distinct ARGs among treatments, with significantly more observed ARGs in the TWW-irrigated enrichments relative to the litter-amended enrichments, which were significantly higher than the freshwater-irrigated enrichments (Fig. 3B,C).

### Co-occurrence of clinically important under-the-radar bacterial taxa and ARGs

Co-occurrence analysis was used to correlate between selected pathogen-harboring genera, ARGs, and MGEs. We included fecal coliforms as a “gold standard” bacterial indicator, together with *Salmonella*, *Pseudomonas*, *Acinetobacter* and *Aeromonas*, which can be clinically-important, are frequently abundant in the environment, and were also previously linked to inter-phyla transmission of mobile ARGs^10,25^. We also included clinically relevant ARGs and MGEs. The *Escherichia-Shigella* genus was strongly linked to extended-spectrum-β-lactamase (ESBL) encoding genes (*i.e. bla*_CTX-M_, *bla*_TEM_, and *bla*_SHV_), and to genes associated with fluoroquinolone and sulfonamide resistance. Several ARGs co-occurred in *Escherichia-Shigella* and *Klebsiella*, including the ESBL encoding gene *bla*_TEM-7_. No co-occurrence was observed between *Aeromonas* and the targeted β-lactamases, but interestingly, *Aeromonas* was linked to the same mobile colistin resistance gene MCR-7 as *Pseudomonas*, and this co-occurrence correlates to the fact that both genera were induced in the aerobic enrichments. *Enterobacter* was not connected to the primary *Aeromonas-Escherichia-Shigella*-*Klebsiella* network but did share a few common ARGs with *Salmonella* and *Acinetobacter*. Several transposases and insertion sequences (IS), including *tnpA*, *IS*1, and *IS*1394, were found to be linked to multiple ARGs (Fig. 4).

**Fig. 4.**
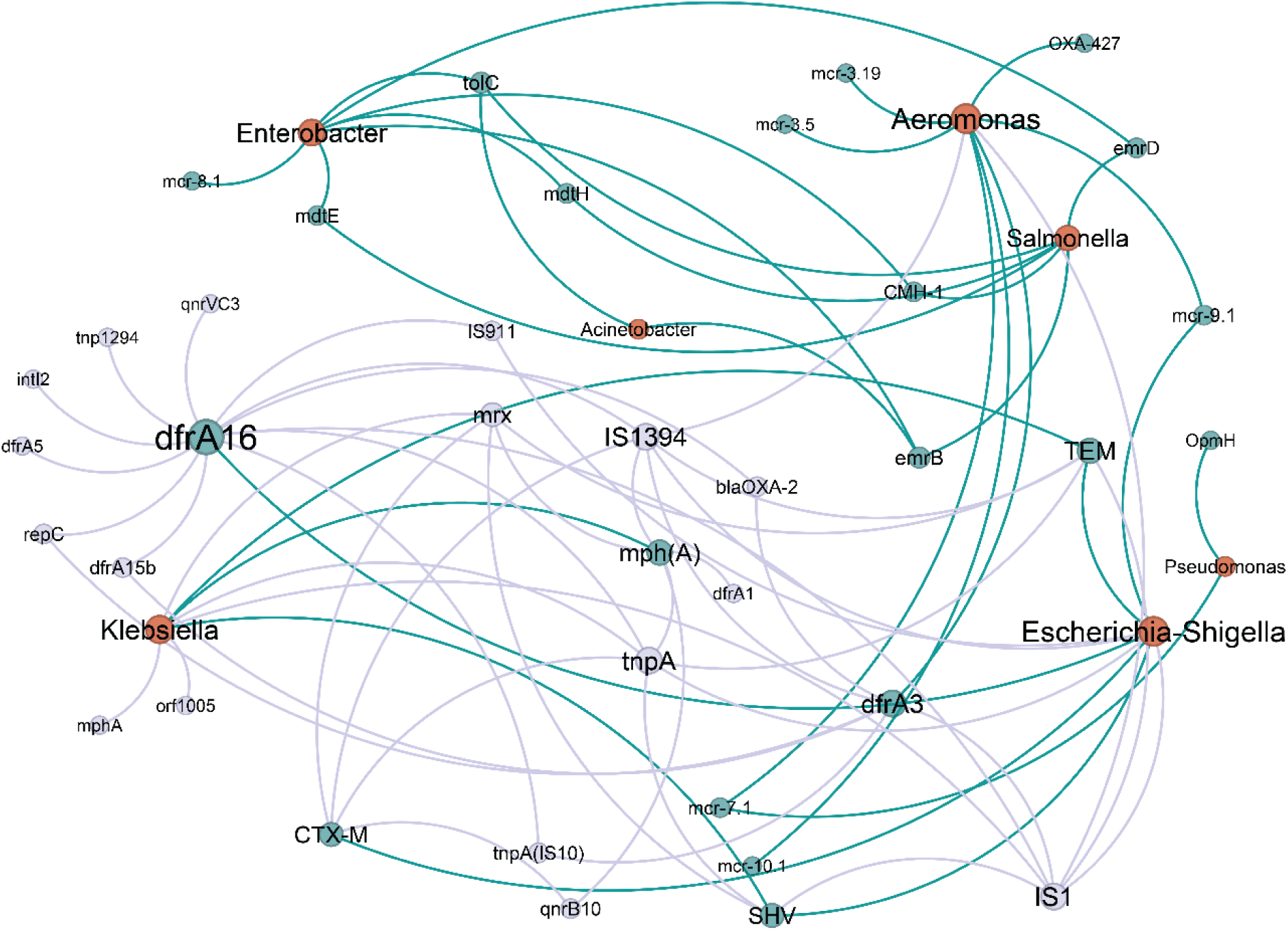
Network analysis showing correlations between rank I ARGs, MGEs, and bacterial genera extrapolated from metagenomic analysis of rhizosphere and phyllosphere enrichments. Nodes represent ARG subtypes, MGEs, and bacterial genera, and edges indicate strong (Spearman rho > 0.7) and significant (padj < 0.01) pairwise correlations. The size of each node is proportional to the number of connections (i.e., degree). Red- Bacteria, turquoise- ARGs, gray- MGEs. ARG ranking followed the pipeline outlined by Zhang et al.^26^, with rank data sourced from Xu et al.^27^. Unranked genes were evaluated using gene information available on the CARD database (https://card.mcmaster.ca/).

### Horizontal transfer of ARGs from under-the-radar clinically-relevant isolates to human pathogen indicators

Co-occurrence of clinically relevant bacteria, ARGs, and MGEs suggested that “under the radar” ARB isolates from anthropogenic enrichments could horizontally transfer ARGs to sensitive strains. ERIC PCR analysis of these isolates revealed a small number of strains that dominated the anthropogenically impacted enriched samples, suggesting that enrichment causes clonal expansion. We established a collection of 9 cefotaxime-resistant *Enterobacteriaceae* strains (6 *E. coli*, 2 *K. pneumoniae*, and 1 *Enterobacter cloacae*) isolated from the enriched anthropogenically affected rhizosphere and phyllosphere samples. We then evaluated the capacity of these strains to horizontally transfer cefotaxime resistance to sensitive pathogen indicators using a custom mating platform. Cefotaxime resistance transfer was observed in two out of the 9 strains, with calculated HGT frequencies ranging from ∼0.0002% to ∼0.03% (Supplementary Fig. 6). Whole genome sequencing (WGS) of these two strains revealed that both harbored *bla*_CTX-M_, as well as additional β-lactamases, and WGS of the transconjugant (PL-W2-608) revealed the acquisition of a plasmid from *E. coli* donor strain C-PL-W2 that harbored six ARGs: *aadA5*, *dfrA17*, *mph(A)*, *qnrS1*, and *sul1* in addition to *bla*_CTX-M-15_ (Fig. 5A). This genotype correlated to phenotypic acquisition of ampicillin, ceftriaxone, chloramphenicol, and trimethoprim-sulfamethoxazole, along with intermediate resistance to ciprofloxacin (Fig. 5B). Surprisingly, the same plasmid was transferred from the second *E. coli* donor Ecl-PL-W1, which also contained several plasmid-associated ARGs on its chromosome. The chromosomes of both donors also contained additional ARGs and several virulence genes associated with extraintestinal pathogenic *E. coli* (ExPEC) and urinary tract infections in humans and animals^28–31^. Non-conjugative *E. coli* strains, and *K. pneumoniae* and *E. cloacae* isolates, did not possess crucial virulence genes, but were all found to contain various ARGs, and displayed MDR phenotypes in susceptibility assays using various antibiotics.

**Fig. 5.**
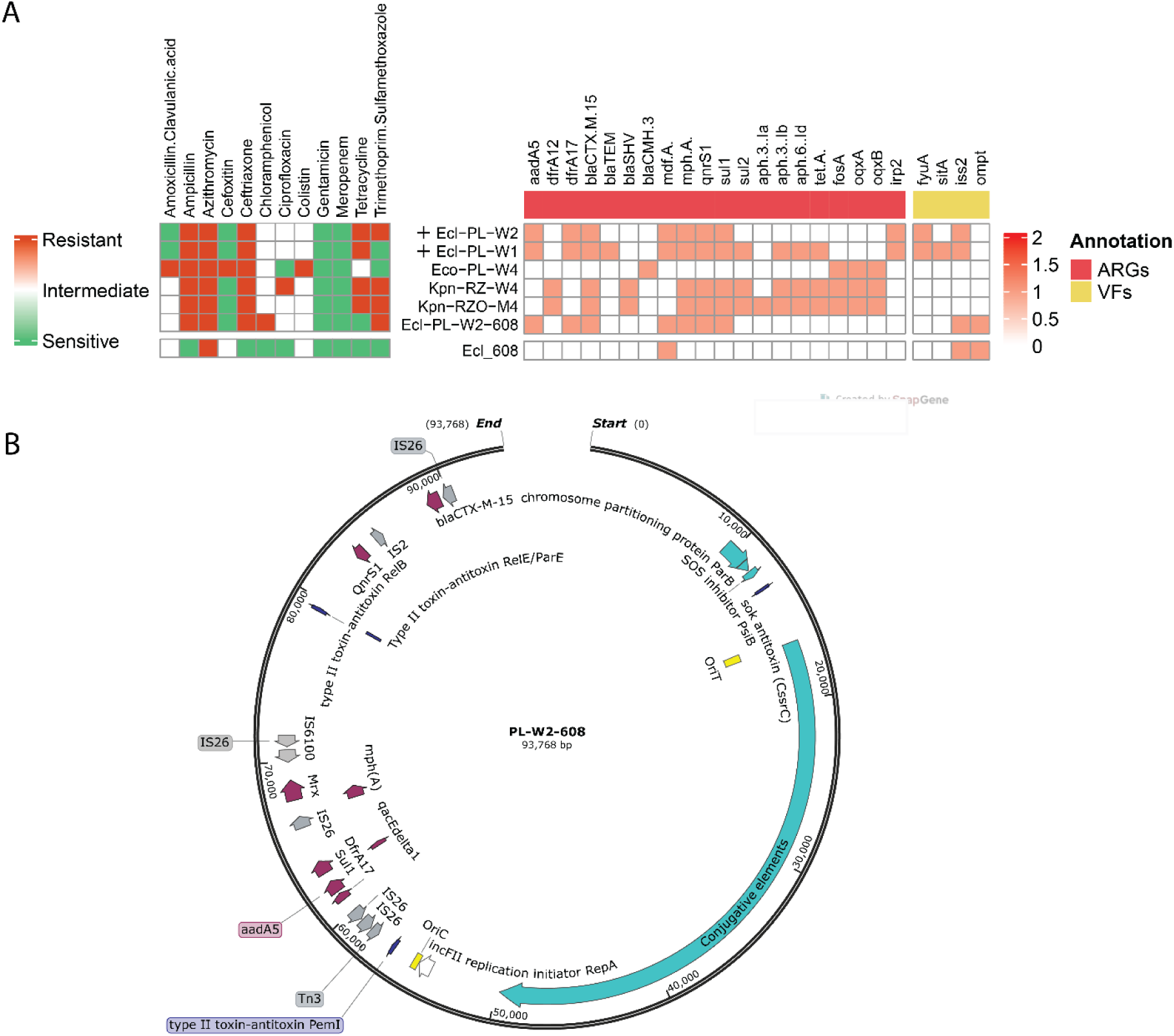
Genotyping and antibiotic susceptibility of Enterobacterial strains isolated from litter-amended and TWW-irrigated enrichments. **A** Antibiotic susceptibility, ARGs, and virulence genes of enterobacterial isolates. **B** Genetic map of the MDR-associated conjugative plasmid detected in this study. The presented plasmid was found in two different *E. coli* donor strains from the wastewater-irrigated rhizosphere and phylosphere enrichments shown in (A). Ecl- *E.coli*, Eco- *Enterobacter Cloacae*, Kpn- *Klebsiella pneumoniae*, + indicates that the plasmid is conjugative.

Overall, the mating experiments demonstrated that cefotaxime-resistant *E. coli* strains that proliferated in anthropogenic-affected, enriched samples have the potential to transfer ESBLs to sensitive strains. These donors, and the other coliform isolates exhibited MDR phenotypes with corresponding resistance genes, and were related to nosocomial pathogenic strains.

## Discussion

The results presented in this study underline the fact that direct assessment of pathogen and AMR markers in anthropogenically-impacted agroecosystems using cultivation-based^6^ or molecular^7^ methods can neglect clinically important markers that are present below the LOD. This is critical considering that unlike chemical contaminants, bacteria can multiply and rapidly proliferate under favorable conditions, potentially becoming hazardous. Brain Heart Infusion (BHI) broth was previously reported to enrich bacteria in a manner representative of the gut microbiome^32,33^. Enrichment in BHI stimulated growth of clinically-relevant bacterial genera, proportionately to the level of anthropogenic exposure (TWW-irrigation > litter amendment > freshwater irrigation), coinciding with previous studies that demonstrated that copiotrophic-stimulating media induces pathogen-associated populations in environmental samples that are otherwise overlooked by direct analyses^13,14^.

As previously demonstrated^34^, most of the bacterial taxa detected in the direct molecular analyses were characteristic of soil and rhizosphere ecosystems, with almost no detected clinically relevant pathogen or ARG indicators, despite their profuse presence in the low-quality irrigation water and un-stabilized litter amended to the soil^35,36^. We attribute this phenomenon to ecological barriers facilitated by a robust soil microbiome that prevents colonization by invaders through nutrient and spatial competition and antibiosis^37^. The native phyllosphere microbiomes and resistomes were considerably more variable than those of the rhizosphere. This stochasticity may be attributed to the high ecological variation (e.g. UV radiation, temperature, dust, insects, etc.) of the phyllosphere relative to the root environment, or to the lower stability of the phyllosphere in modulating its local microbiome relative to the rhizosphere^38^.

In lettuce rhizosphere and phyllosphere samples, enrichment resulted in a decrease in α-diversity compared to native samples, due to physicochemical conditions that stimulate growth of the copiotrophic fraction of the original community, a phenomenon previously described as “enrichment bias”^39^. Under aerobic and anaerobic enrichments, this bias stimulated bacterial taxa within the Bacillota and Pseudomonadota phyla, at the expense of oligotrophic taxa that are more characteristic of natural environments^40^, similar to previously reported findings^14^. From an ecological perspective, this bias is highly detrimental, but from a public health/risk assessment perspective, it enabled us to identify pathogen-associated taxonomic and ARG indicators, with significant correlations between treatment (freshwater irrigation, litter-amendment, and TWW irrigation) and the bacterial and ARG profiles. While this “anthropogenic footprint” was much more deterministic in the enriched rhizosphere samples, it was still significant in the phyllosphere samples, despite the fact that the lettuce plants in the present study were drip irrigated, and there was no direct contact between the irrigation water or amended litter and the leaves. This suggests that AMR determinants may have been transported from the rhizosphere to upper plant compartments, a phenomenon that was previously described when pathogens inoculated onto roots were capable of trafficking to the phyllosphere in edible plants^41,42^. Enrichment cultures with and without oxygen resulted in similar profiles, suggesting that many of the enriched taxa are facultative anaerobes, e.g. members of the *Enterobacteriaceae* and *Bacillaceae* families. Three exceptions were *Paraclostridium*, which was more abundant in the anaerobic enrichment, and *Pseudomonas* and *Acinetobacter*, which proliferated more in the aerobic enrichments, accordingly to their respective atmospheric growth requirement. HTqPCR analysis as well as MALDI-MS of strains isolated from enriched aerobic litter-amended phyllosphere revealed the presence of *A. baumannii* (Supplementary table 2), an opportunistic pathogen with considerable medical importance^20^. Several *Pseudomonas* spp. are also known as opportunistic pathogens^43^, and while *Paraclostridium* is profuse in soil environments and generally not considered hazardous, recent studies indicate that some strains can be pathogenic^44^. Thus, both aerobic and anaerobic enrichment should be considered in future studies that assess microbial and AMR risks in anthropogenically impacted environments.

Combined evaluation of culture-based, amplicon sequencing and shotgun metagenomic analyses highlighted the capacity of enrichment to identify pathogen-associated genera including *Escherichia-Shigella*, *Klebsiella*, *Enterobacter*, *Salmonella*and *Enterococcus* in anthropogenically impacted samples. This was especially evident in the capacity to enrich cefotaxime-resistant coliforms and *E. coli* harboring ESBL-encoding genes and a myriad of other ARGs. Under a “One Health” diagnostic approach, ESBL-producing *E. coli* were suggested as gold standard indicators for AMR risk assessment of animal husbandry facilities and the environment^16–18^. The fact that anthropogenically-impacted phyllosphere coliforms were detectable using culture-based methods but not by 16S amplicon sequencing, while metagenomic analysis identified several pathogens that may be difficult to cultivate, highlights the advantages of combining molecular tools and cultivation based approaches for environmental surveillance^6^. The higher abundance of clinically relevant bacteria in the enriched anthropogenically affected rhizosphere and phyllosphere samples strongly correlated to increased richness of MGEs and ARGs, including genes conferring resistance to an array of clinically important antibiotics. These included ESBL-encoding genes, such as *bla*_TEM_, *bla*_SHV_, *bla*_CTX-M_, and *bla*_VIM_ that co-occurred with pathogen-associated genera. These results were supported by the presence of multiple ARGs including ESBL-encoding genes in the genomes of strains isolated from litter-amended and TWW-irrigated rhizosphere enrichments. The detection of *E. coli*isolates from enriched anthropogenically impacted samples capable of horizontally transferring up to 6 different ARGs to a sensitive *E. coli* recipient may be a particular concern to public health. Moreover, the *E. coli* isolates capable of transferring cefotaxime-resistance were characterized as ST10 and ST1598 and contained the Yersinia high pathogenicity island of virulence factors, which are both common genotyping features in uropathogenic *E. coli*, a prevalent pathotype of this species. Thus, our approach may not only facilitate detection of total and resistant bacterial pathogen indicators, but can also reveal the presence of specific strains of considerable medical importance.

Additionally, using different enrichments with various culture media can help reduce the variability often seen in highly stochastic environmental samples like the phyllosphere, making it easier to analyze the desired fraction of the native community. Here, enrichment helped detect AMR determinants in anthropogenically-affected phyllosphere, demonstrating how intense anthropogenic pressure on the rhizosphere can lead to the transfer of AMR determinants to the upper parts of plants, potentially ending up in edible tissues and posing a public health concern. Various platforms have been developed to rank the risks of ARGs^26,45^, and microbial source tracking (MST) and quantitative microbial risk assessment (QMRA) have been applied (separately and in tandem) to evaluate risks of anthropogenically-impacted environments^46^. Certain QMRA methods not only estimate hazards linked to resistant pathogens, but also integrate risks associated with horizontal transfer of specific ARGs^47^. Nonetheless, assessing risks associated with AMR in anthropogenically impacted environments is extremely challenging considering the dynamics of AMR and the high complexity of environmental microbiomes. Based on our study, this challenge is further exacerbated by the fact that many pathogens and ARGs that are traditionally targeted in QMRA are present below LOD in anthropogenically impacted ecosystems. We believe that the novel enrichment approach depicted in this study (integrating culture-based, molecular and HGT assays) can be fundamental in providing “under the radar” empirical data that can significantly enhance QMRA models and have a groundbreaking impact on AMR risk assessment that can be translated to better policies. This is critical considering the global spread of AMR, and the increasing application of manure and wastewater for agriculture due to growing global populations and water scarcity due to climate change.

## Methods

### Lettuce sampling, enrichment cultures, and gDNA extraction

Lettuce seedlings were planted in mid-August in the Maayan Zvi wastewater treatment plant located in northern Israel. Each pot represented a biological replicate and contained four plants. The treatments applied in this study included freshwater irrigation, amendment with poultry litter from broilers (5 cubic meters per dunam), and irrigation with low-quality secondary wastewater effluent (applied for 15 min, three times a day, at 2 liters per hour), with four replicates per treatment. All pots were chemically fertilized prior to planting to minimize the impact of the anthropogenic treatments on plant growth. After 45 days of growth, the plants were uprooted, and the leaves (phyllosphere) and roots (rhizosphere) were separated. The root samples were shaken over a weighting plate to collect the soil attached to the roots and was transferred into 50 ml falcon tubes. The leaves were washed vigorously using a freshwater hose, and 30 grams of leaves were weighed and placed in a sterile stomacher bag. All samples were kept in cooling boxes with ice packs until processing. In the lab, 0.85% NaCl sterile saline (w/v) was added to the phyllosphere stomacher bags, which were then processed in a stomacher for 2 minutes, and the liquid was collected from the filtered side into 50 ml tubes. For the rhizosphere samples, 5 grams were collected from each sample, and 30 ml of 0.85% saline was added along with glass beads. The mixture was vortexed at maximum speed for 1 minute, and then horizontally shaken for 20 minutes at 500 rpm. Lysates were centrifuged for 10 minutes at 600 rpm at 4°C, and 20 ml of the supernatant were transferred to a new tube. This process was repeated once more resulting in a final supernatant volume of 12 ml. Copiotrophic enrichment cultures were prepared by inoculating 100 µl of the clear phyllosphere and rhizosphere lysates into pre-made anaerobic Hungate tubes (sparged with N2) and aerobic culture tubes containing BHI medium, which has shown good recovery of gut microbial communities^32,33^. Anaerobic inoculation was done using a sterile needle. Additionally, 1 ml of phyllosphere and rhizosphere lysates was added to 750µl of 50% glycerol for further culture-based analyses. Enrichments were grown for 16 hours at 180 rpm and 37°C, and glycerol stocks were prepared in the same manner. Subsequently, Phyllosphere lysates were centrifuged at 5000 rpm and 4°C for 10 minutes, and the supernatants were discarded. The pellets were stored at −80°C for further gDNA extraction. At the time of extraction, all cultures were centrifuged for 1 minute at 5000 rpm, and supernatant was discarded. Pellets from enrichment cultures and phyllosphere lysates were extracted using the MagAttract High Molecular Weight (HMW) kit (Qiagen). gDNA from Rhizosphere samples was extracted using the Powersoil kit (Qiagen). gDNA quality was assessed by measuring 260/280 and 230/260 ratios with the Nanodrop system 2000c Spectrophotometer (Thermo Fisher Scientific) and quantified using a Qubit fluorometer (Qubit 2.0, Invitrogen).

### Fecal coliform quantification

Glycerol stocks were completely thawed, serially diluted, and then plated on Chromocult coliform agar (CCA, Merck, Darmstadt, Germany) using the drop method. The plates were incubated for 16 hours at 44°C to select for fecal coliforms, and the colony forming units (CFU) were determined by counting the colonies and multiplying by the appropriate dilution factor.

### Fecal coliform isolation and identification

Cefotaxime resistant coliforms were isolated on CCA plates supplemented with 4 mg/L cefotaxime. Bacteria were selected based on colony color on the CCA agar (blue for *Escherichia-Shigella*, pink for other coliforms), and identified by MALDI-TOF-MS (score >2.0) using an Autoflex system, following the direct colony method (Bruker Daltonics, Billerica, MA, USA). To avoid multiple isolations of the same strain, isolates were analyzed using the enterobacterial repetitive intergenic consensus polymerase chain reaction (ERIC-PCR) method for genotyping. Finally, selected isolates were stored in glycerol stocks at −80 C°.

### Quantitative PCR

The composition and relative abundance of selected ARGs and MGEs in litter samples were analyzed using a custom HTqPCR array, specifically targeting 53 clinically relevant ARGs and MGEs selected from a previously described set of primers^48^. Enrichments (n=48 of anaerobic and aerobic cultures) were transferred to the Key Lab of Urban Environment and Health at the Institute of Urban Environment, Chinese Academy of Sciences (Xiamen, China), where HT-qPCR was performed using the Wafergen Smart Chip Real-time PCR system as previously described by Muurinen et al^49^.

### 16S rRNA gene sequencing and data analysis

Native samples and enrichments microbiota were further profiled by sequencing the V4 region of the 16S rRNA gene for all DNA samples. Extracted DNA was amplified using the CS1-515F_new and CS2-806R_new barcoded primers, which target the V4 region of the 16S rRNA gene to generate sequencer-ready libraries as previously described^50^. The barcoded libraries were pooled and sequenced on an Illumina MiniSeq platform with a 150-bp paired-end strategy, employing V3 chemistry for 16S genes at the Rush University DNA Services Facility, Chicago. Due to high plastid contamination, phyllosphere samples were sequences four times with a plastid blocker in addition to the initial run. The generated paired-end sequences were merged using PEAR^51^ and processed using QIIME2 v2019.7^52^. The sequences underwent quality filtering employing the DADA2 algorithm^53^, which resolves amplicon sequence errors to generate amplicon sequence variants (ASVs). Taxonomic assignment was performed using the QIIME2 q2-feature-classifier with the SILVA 132 rRNA database^54^. Samples with low sequencing depth such as negative controls (< 1000 reads) were excluded from further analysis, as were ASVs corresponding to plastids or mitochondria. A rooted phylogenetic tree was built using FastTree (v2.1.10)^55^, based on the generated ASVs. Raw count tables, taxonomic assignments, and the rooted phylogenetic tree were then exported for further analyses and visualization in R.

### Shotgun metagenome sequencing and analysis

Library preparation and sequencing of enrichment samples were performed at Guangdong Magigene Biotechnology Co. Ltd. (Guangzhou, China). Sequencing libraries were generated using ALFA-SEQ DNA Library Prep Kit following manufacturer’s recommendations and index codes were added. The library quality was assessed on the Qubit 4.0 Fluorometer (Life Technologies, Grand Island, NY) and Qsep400 High-Throughput Nucleic Acid Protein Analysis system (Houze Biological Technology Co, Hangzhou, China) system. The library was then sequenced on an Illumina NovaSeq 6000 platform, generating 150 bp paired-end reads. Read quality was evaluated using the FastQC tool^56^ and adjusted with Trimmomatic software^57^. De novo genome assembly, binning, and MAGS construction of processed reads were achieved using the MetaWRAP pipeline^58^. Taxonomic classification was performed using kaiju (v1.9.2)^59^ with default parameters and the NCBI BLAST NR reference database was selected^60^. To characterize ARGs, we employed ARGs-OAP (v3.2.3)^61^, and used the unnormalized count subtype output for further analyses. Open reading frame (ORF) prediction was performed on scaffolds using Prodigal^62^. ORFs were subjected to BLASTn (v2.6.0+ maximum e-value 1×e^−10^) search against the INTEGRALL database^63^ for MGEs annotation. Identification of ARGs that were differentially abundant in TWW irrigated, litter amended and freshwater irrigated enrichments was achieved using DESeq2^64^, following alignment to the ResFinder database^24^.

### Co-occurrence analysis

Network analysis was performed to reveal the underlying associations between ARGs, MGEs, and microbial taxa. To create a count table from the MGEs analysis output, a nonredundant set of ORFs was obtained through CD-HIT (v4.8.1)^65^ applying a 95% identity threshold. Reverse mapping took place using bowtie2 (v2.3.5.1)^66^, and Samtools (v1.9)^67^ was used to sort the read mappings. A correlation matrix was constructed by calculating all possible pairwise Spearman correlation coefficients (rho) among ARG subtypes and bacterial genera, ARG subtypes and MGEs, and MGEs and bacterial genera based on the abundance data from the metagenomic analysis using the Hmisc R package (v5.1-1)^68^. Features fewer than 50 reads across all samples were removed prior to the analysis. ARG ranking was determined based on the pipeline described by Zhang et al^26^. Only statistically strong (rho > 0.70) and significant correlations (padj < 0.01) were retained for this study. Finally, the Gephi software package (v0.10.1) was used to visualize the correlation network.

### Assessment of horizontal gene transfer

Based on ERIC-PCR patterns and MALDI results, 29 strains were selected for the next mating experiment. The rifampicin- and spectinomycin-resistant GFP-tagged *E. coli* BW25133 GFP #608 was used as the recipient strain. All donor strains were cultivated in triplicates in 2 ml Luria Broth (LB) supplemented with cefotaxime (4 mg/L) and incubated overnight at 37°C with shaking at 160 rpm. The recipient strain was incubated in 5 ml LB supplemented with rifampicin (50 mg/L). All bacterial combinations were washed twice in a corresponding volume of LB without antibiotics. Subsequently, 250 µl of the recipient culture and 250 µl of each donor culture were mixed together in 500 µl of LB in 1.5 ml Eppendorf tubes and incubated for 24 hours with gentle shaking. After incubation, 100 µl aliquots were taken from each tube, mixed with 1 ml of 0.85% NaCl (w/v) saline solution, and serially diluted. 100 µl of appropriate dilution was plated on LB agar plates supplemented with cefotaxime (4 mg./L) and rifampicin (50 mg/L).

In parallel, 100 µl from the original recipient culture was diluted and plated on LB agar plates supplemented with rifampicin (50 mg/L). After 24 h of incubation at 37°C, the CFU of recipient and transconjugants were calculated. Transconjugants were confirmed by GFP fluorescence using a fluorescence binocular. HGT frequency was calculated using the formula: Transfer frequency= (CFU ml^−1^ transconjugants)⁄(CFU ml^−1^ recipients).

### Long and short reads Whole Genome Sequencing (WGS) and analysis

DNA library preparations were generated for short-read whole genome sequencing following the Hackflex protocol^69^, a modified version of the Illumina DNA preparation. Sequencing was performed on an Illumina NovaSeq 6000 platform. Short-read whole genome sequences were assembled using shovill (v1.0.9) (https://github.com/tseemann/shovill). Genotyping was performed with ABRicate (https://github.com/tseemann/abricate) using the following databases for each set of detected genes: ResFinder^24^, CARD^70^, PlasmidFinder^71^, ISFinder^72^, VFDB^73^, and the *E. coli* customDB (https://github.com/maxlcummins/E_coli_customDB). Sequencing typing, and any potential plasmid subtyping were performed with MLST (https://github.com/tseemann/mlst). Selected isolates were subjected to further long-read Nanopore sequencing by Plasmidsaurus (Oregon, US). Hybrid assembly was performed using Unicycler (v0.5.0)^74^, and genes were annotated with Bakta (v1.9.2)^75^.

### Antimicrobial susceptibility testing

Eight isolates were tested for antimicrobial susceptibility to 13 antibiotics (Amoxicillin/ Clavulanic acid, Ampicillin, Azithromycin, Cefoxitin, Ceftriaxone, Chloramphenicol, Ciprofloxacin, Gentamicin, Meropenem, Nalidixic acid, Sulfisoxazole, Tetracycline, and Trimethoprim/Sulfamethoxazole) with the CMV4AGNF AST Plate (Sensititre, TREK diagnostic systems, Thermo scientific), following the manufacturer’s instructions. Plates were read after overnight incubation in 37°C with the Vizion apparatus (Thermo scientific). Minimal inhibitory concentrations were interpreted following the CLSI breakpoints^76^.

### Statistical analysis and visualization

Significant differences were assessed using several methods detailed in the main text and figure legends. Diversity indices for alpha and beta diversity were analyzed using the Pairwise Wilcoxon test and PERMANOVA, respectively, with the vegan package in R (v2.6.4)^77^. Wilcoxon test for multiple comparisons of HTqPCR and CFU results were calculated using JMP17 software. Differentially abundant taxa and genes were detected using the DESeq2 package (v1.38.3)^64^, and heatmaps were generated using the pheatmap package in R (v1.0.12)^78^. CFU plots were visualized with GraphPad Prism (v8.4.3), and Venn diagrams were generated with the online tool https://bioinformatics.psb.ugent.be/webtools/Venn/.

