## Supplementary information for "Occurrence of “under-the-radar” antibiotic resistance in anthropogenically affected produce"

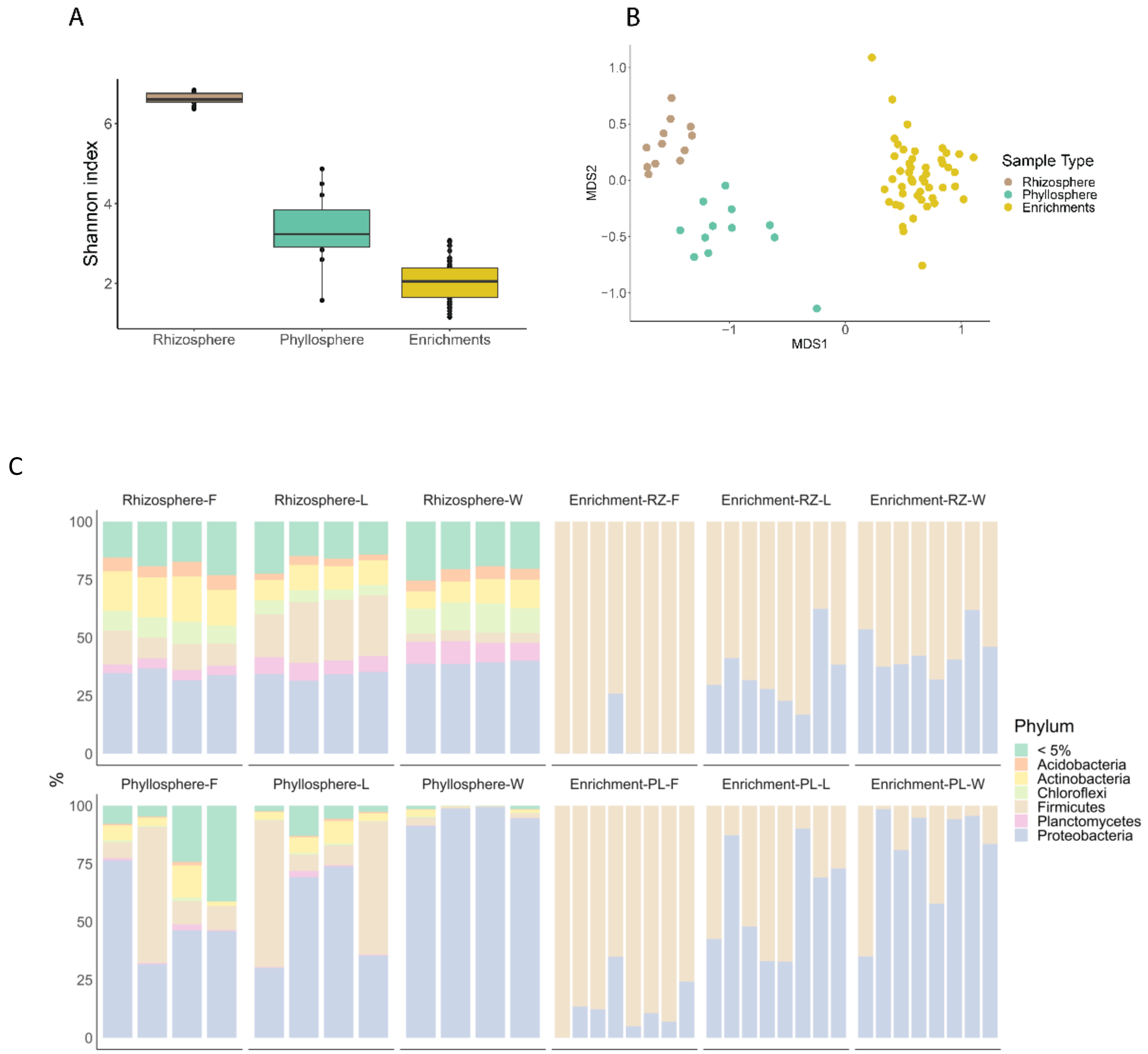


**Supplementary Fig. 1**. Impact of treatment and enrichment on bacterial diversity and community composition based on 16 rRNA gene amplicon sequencing data. **A** α-diversity based on the Shannon diversity index of native and enriched samples (Wilcoxon test $p<0.00001$. Figure 1A). **B** β-diversity of rhizosphere, phyllosphere, and enrichments bacterial communities using Bray–Curtis distance matrix and visualized by NMDS plot analysis (PERMANOVA, p.value < 0.05) **C** Bacterial community composition (phylum-level) of source and enriched rhizosphere and phyllosphere samples. RZ= rhizosphere, PL= phyllosphere, F= freshwater, L= litter, W= TWW. <5% indicates families whose relative abundance represents < 5% of the total characterized bacterial community.

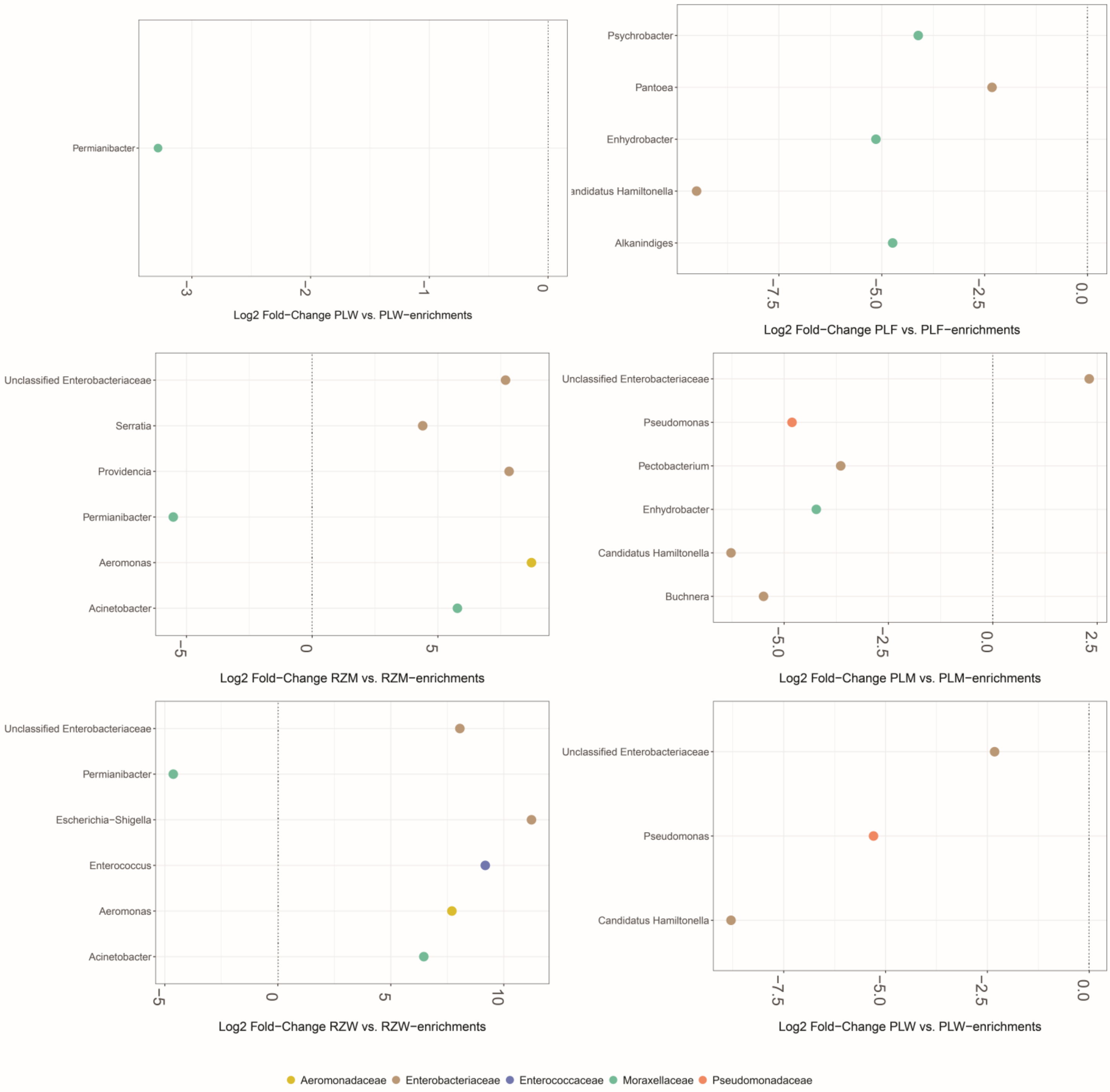


**Supplementary Fig. 2**. Differential abundance of genera from pathogen-associated families between enrichment and native samples. Positive log2 fold change values indicate higher abundance in enrichments, while negative values indicate higher abundance in native samples. (DESeq2 analysis: log2FoldChange > 1, padj < 0.01).

| Factor | baseMean | log2FoldChange | lfcSE | stat | pvalue | padj | *Genus* |
|---|---|---|---|---|---|---|---|
| PLM | 14344.84 | 6.035623 | 1.147809 | -5.25839 | 1.45E-07 | 9.95E-06 | *Bacillus* |
| PLM | 27173.42 | 4.497438 | 1.401127 | -3.20987 | 0.001328 | 0.006738 | *Unclassified Enterobacteriaceae* |
| PLW | 1100.901 | 3.095895 | 0.991174 | -3.12346 | 0.001787 | 0.008749 | *Bacillus* |
| RZM | 229.7507 | 26.35921 | 3.091026 | -8.52766 | 1.49E-17 | 1.18E-16 | *Providencia* |
| RZM | 2446.945 | 10.7966 | 2.665583 | -4.05037 | 5.11E-05 | 9.94E-05 | *Aeromonas* |
| RZM | 12443.98 | 9.910753 | 1.384442 | -7.15866 | 8.15E-13 | 3.29E-12 | *Unclassified Enterobacteriaceae* |
| RZM | 1715.905 | 8.079696 | 1.585899 | -5.09471 | 3.49E-07 | 8.1E-07 | *Acinetobacter* |
| RZM | 9.974079 | 7.60409 | 2.461528 | -3.08917 | 0.002007 | 0.003345 | *Serratia* |
| RZM | 671.0401 | 7.070627 | 1.413969 | -5.00055 | 5.72E-07 | 1.31E-06 | *Clostridium sensu stricto 1* |
| RZM | 32146.54 | 5.130042 | 0.561444 | -9.13724 | 6.41E-20 | 7.15E-19 | *Bacillus* |
| RZM | 1056.85 | 2.304 | 0.701803 | -3.28297 | 0.001027 | 0.001755 | *Lysinibacillus* |
| RZW | 1688.045 | 28.18871 | 3.078817 | -9.1557 | 5.4E-20 | 3.44E-19 | *Paraclostridium* |
| RZW | 1521.663 | 13.8925 | 1.143573 | -12.1483 | 5.86E-34 | 1.49E-32 | *Escherichia-Shigella* |
| RZW | 300.1966 | 11.55189 | 1.167782 | -9.89216 | 4.5E-23 | 4.19E-22 | *Enterococcus* |
| RZW | 2913.691 | 8.253645 | 0.786566 | -10.4933 | 9.28E-26 | 1.13E-24 | *Unclassified Enterobacteriaceae* |
| RZW | 4155.529 | 7.841534 | 0.720064 | -10.8901 | 1.29E-27 | 1.84E-26 | *Aeromonas* |
| RZW | 360.2252 | 6.724341 | 2.187663 | -3.07376 | 0.002114 | 0.003321 | *Acinetobacter* |
| RZW | 8016.005 | 5.51849 | 0.881317 | -6.26164 | 3.81E-10 | 9.62E-10 | *Bacillus* |
| RZW | 2794.81 | 4.703947 | 0.829026 | -5.67407 | 1.39E-08 | 3.19E-08 | *Clostridium sensu stricto 1* |
| Anaerobic | 892.8958 | -7.63186 | 0.874408 | 8.728027 | 2.59E-18 | 3.81E-15 | *Paraclostridium* |
| Anaerobic | 45.45833 | -6.49051 | 0.7619 | 8.518847 | 1.61E-17 | 1.18E-14 | *Terrisporobacter* |
| Aerobic | 3052.625 | 5.759358 | 0.856405 | -6.72504 | 1.76E-11 | 8.60E-09 | *Acinetobacter* |
| Aerobic | 150.6875 | 3.72663 | 0.713726 | -5.22138 | 1.78E-07 | 6.52E-05 | *Pseudomonas* |
| Anaerobic | 4.729167 | -3.08037 | 0.629359 | 4.894462 | 9.86E-07 | 0.00029 | *Morganella* |
| Aerobic | 4.0625 | 2.832888 | 0.623446 | -4.54392 | 5.52E-06 | 0.001352 | *Providencia* |
| Enrichments | 3508.503 | 6.737762 | 1.43169 | -4.70616 | 2.52E-06 | 1.04E-05 | *Lactococcus* |
| Enrichments | 9710.694 | 3.227303 | 0.747125 | -4.31963 | 1.56E-05 | 5.48E-05 | *Bacillus* |

**Supplementary table 1**. Influence of enrichment, oxidation levels, and treatment factors on bacterial genera abundance. RZ- rhizosphere; PL- phyllosphere; L-litter; W- wastewater; FC-fold change; UC- unclassified (Deseq2 p.adj<0.01 log2FoldChange>1). FC-fold change.

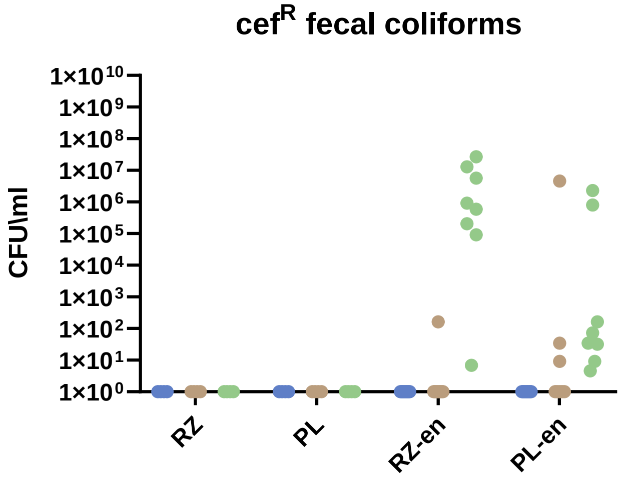


**Supplementary Fig. 3**. CFU/ml of cefotaxime resistant of total cefotaxime resistant fecal coliforms (Wilcoxon pairwise test p.value <0.0001)

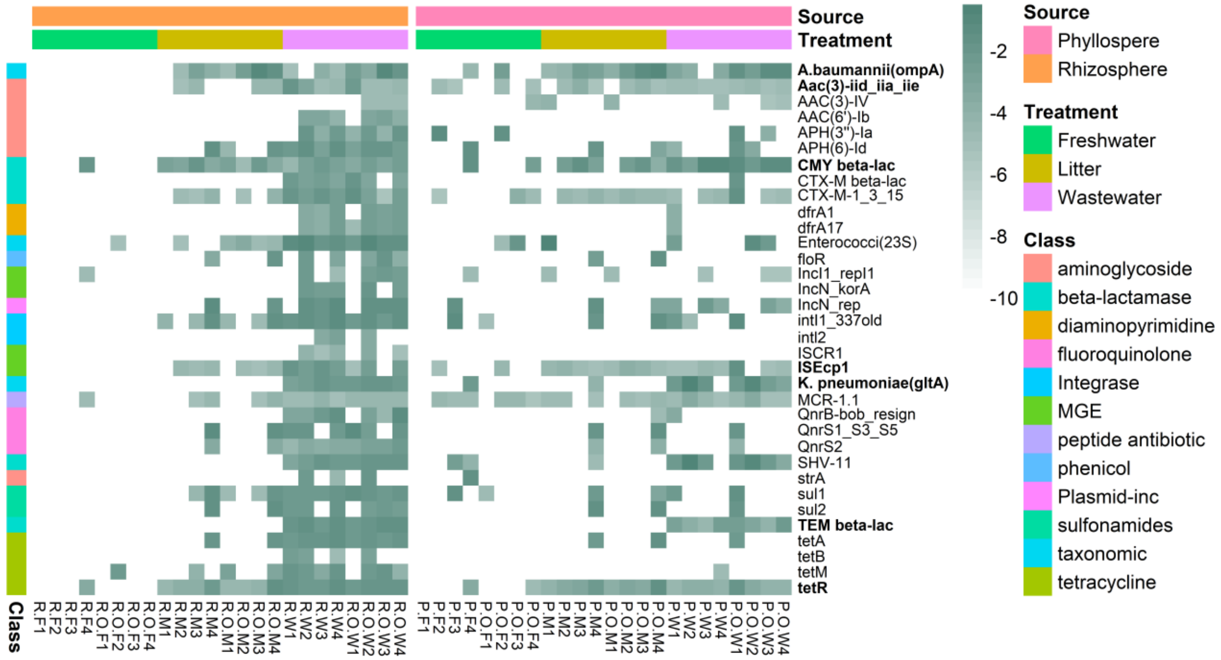


**Supplementary Fig. 4**. HT-qPCR analysis of enriched rhizosphere and phyllosphere samples targeting clinically-relevant ARGs and taxa, showing gene markers with statistically significant differences between the three treatments. Bolded markers indicate that they were significant in both rhizosphere and phyllosphere anthropogenically-associated enrichments (Wilcoxon Test p.value<0.05)

A

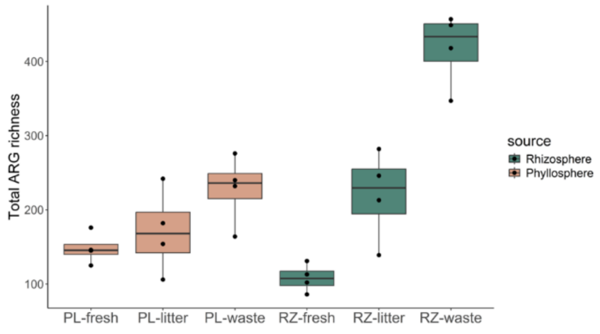


B

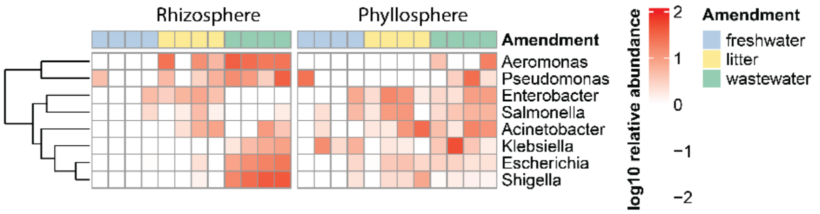


**Supplementary Fig. 5.** ARGs and taxa abundance and diversity based on metagenomic data. **A** ARGs richness of freshwater-irrigated, litter-amended, and TWW-irrigated enriched rhizosphere and phyllosphere samples (Wilcoxon test p<0.05). **B** log10 relative abundance of clinically-relevant genera in enriched samples (Deseq2 p.adj<0.01). RZ- rhizosphere; PL- phyllosphere; fresh- freshwater; waste- wastewater.

| **Isolate** | **Origin** | **Oxidation** | **Detected species** | **Score** |
|---|---|---|---|---|
| C-PLO-M4-1 | Litter-phyllosphere | Aerobic | *K.pneumoniae* | 2.32 |
| C-PLO-M3-5 | Litter-phyllosphere | Aerobic | *K.pneumoniae* | 2.35 |
| C-PLO-M1-1 | Litter-phyllosphere | Aerobic | *K.pneumoniae* | 2.04 |
| C-RZO-M4-4 | Litter-rhizosphere | Aerobic | *K.pneumoniae* | 2.16 |
| C-PL-W4-4 | TWW-phyllosphere | Anaerobic | *E. kobei* | 2.20 |
| C-RZ-W4-5 | TWW- rhizosphere | Anaerobic | *K.pneumoniae* | 2.37 |
| C-PLO-M4-2 | Litter-phyllosphere | Aerobic | *E. asburiae* | 2.17 |
| C-PLO-M4-5 | Litter-phyllosphere | Aerobic | *A. baumannii* | 2.14 |
| C-PLO-M4-3 | Litter-phyllosphere | Aerobic | *E. kobei* | 2.24 |

**Supplementary table 2.** MALDI results of isolates isolated from anthropogenically-associated enrichments

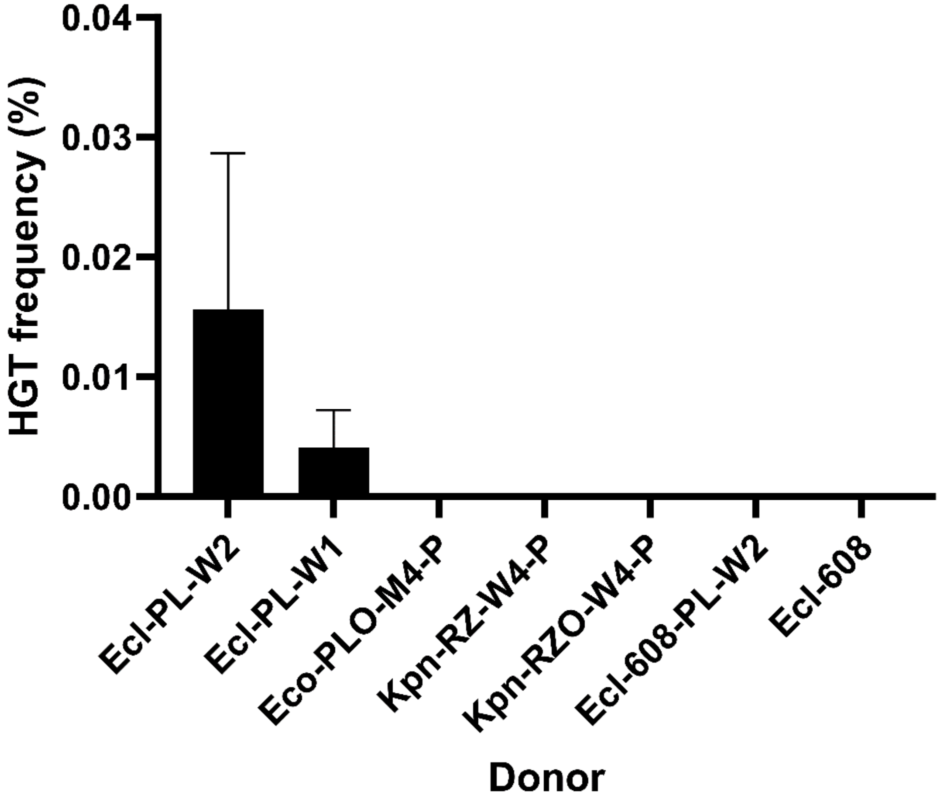


**Supplementary Fig. 6**. HGT of *E.coli 608* by fecal coliform strains isolated from litter- and TWW- associated enrichments represented as the percentage of transconjugants CFU/ml divided by recipients CFU/ml. Ecl- *E.coli*, Eco- *Enterobacter Cloacae*, Kpn- *Klebsiella pneumoniae*.
